# Human iPSC-derived neurons identify West Nile virus neuropathogenesis signatures and reveal strain-dependent neuroattenuation

**DOI:** 10.64898/2026.09.23.753976

**Authors:** Alejandro Matía, Astrid Anaya, Ali Akgul, Anil Singapuri, Sunny Lee, Jinzhao Wang, Ngan Wagoner, Marius Wernig, Thomas C. Südhof, Christopher M. Barker, Michael S. Diamond, Carolina Arias, Ruth Hüttenhain

**Affiliations:** Department of Molecular & Cellular Physiology, Stanford University School of Medicine, Stanford, CA, USA; Chan Zuckerberg Biohub, San Francisco, CA, USA; Department of Medicine, Washington University School of Medicine, St. Louis, MO, USA; Department of Pathology, Microbiology & Immunology, University of California, Davis, CA, USA; Institute for Stem Cell Biology and Regenerative Medicine, Stanford University School of Medicine, Stanford, CA, USA; Department of Pathology, Stanford University School of Medicine, Stanford, CA, USA; Department of Chemical and Systems Biology, Stanford University School of Medicine, Stanford, CA, USA; Howard Hughes Medical Institute, Stanford University School of Medicine, Stanford, CA, USA; Department of Molecular Microbiology, Washington University School of Medicine, St. Louis, MO, USA; Department of Pathology & Immunology, Washington University School of Medicine, St. Louis, MO, USA

## Abstract

West Nile virus (WNV) is one of the leading causes of arthropod-transmitted encephalitis in the world, yet its effects on neurons, the principal target of WNV in the brain, remain poorly defined. Here we establish and use *Ngn2*-induced human iPSC-derived glutamatergic neurons to define the neuronal response to WNV through integrated temporal proteomics and transcriptomics. Temporal multi-omic profiling revealed that infection activated innate immunity and the Unfolded Protein Response while suppressing axonogenesis and synaptic programs. WNV infection also disrupted calcium homeostasis, reduced STMN2 abundance, and promoted cytoplasmic redistribution of TDP-43. Applying this neuronal model to characterize contemporary California isolates, we identified LA24, a lineage 1 strain that replicates efficiently in non-neuronal cell lines but is selectively attenuated in neurons, with reduced viral burden in the brain and attenuated pathogenicity in mice. Together, these findings uncover neuron-specific signatures of WNV pathogenesis and demonstrate that neuronal replication phenotypes can reveal strain-dependent virulence differences not captured by conventional cell lines.

## Introduction

West Nile virus (WNV), a mosquito-borne orthoflavivirus, is the leading cause of arboviral encephalitis in the United States^1–3^. While most infections are asymptomatic, approximately 1 in 140 cases progress to neuroinvasive disease presenting as meningitis, encephalitis or acute flaccid myelitis^4^, with a 13 % case-fatality rate and frequent persistent cognitive, motor and affective deficits among survivors. Despite this burden, no licensed human vaccine or specific antiviral therapy is available for WNV to prevent neuroinvasive disease^5,6^. Systematically mapping how WNV disrupts neuronal molecular programs is a starting point towards the rational design of neuroprotective interventions.

Neurons are the main target of WNV within the brain, and severe disease is associated with neuronal dysfunction, including impaired excitability, synaptic transmission, and axonal transport^7,8^. Yet, defining the molecular changes underlying this neuronal dysfunction has been hindered by the limitations of available model systems: non-neuronal cell lines^9–11^ lack neuronal properties and mount interferon (IFN) responses that differ from those of neurons^12,13^. Conversely, in mouse models and in organoids containing glia and infiltrating leukocytes^14–16^, molecular changes may reflect the effects of viral infection in the context of cellular immune responses, making it difficult to disentangle cell-intrinsic contributions. While primary neuronal cultures partially overcome this limitation by enriching neurons, their limited and variable yield can reduce the depth of multi-omic profiling needed to systematically map infection-induced molecular changes. More recently, human induced pluripotent stem cells (hiPSC)- and embryonic stem cell-derived neurons have provided scalable systems for studying neuron-intrinsic responses. In particular, induced expression of the proneural transcription factor Neurogenin-2 (*Ngn2*) converts hiPSCs within a few weeks into a homogeneous population of excitatory, cortical-like glutamatergic neurons^17^, making this system well suited for multi-omic profiling. hiPSC-derived neurons can also support productive WNV infection^18–20^. However, the molecular response to WNV infection in these neurons has not been systematically mapped, including its timing and extent^21,22^.

Closing this gap requires systematic, temporally resolved measurement of the host proteome and transcriptome during infection in human neurons. During orthoflavivirus infections, molecular changes include activation of the unfolded protein response and endoplasmic-reticulum stress^23^, selective antagonism of IFN-stimulated gene (ISG) induction, and disruption of nucleocytoplasmic transport^24^ which can uncouple transcript accumulation from protein output. Several of these stress axes converge on pathways implicated in chronic neurodegeneration, including calcium dysregulation and loss of the axonal maintenance factor stathmin-2 (STMN2)^25,26^. Epidemiological analyses linking prior viral exposure to subsequent neurodegenerative disease risk raise the question of whether an acute neurotropic infection recapitulates these same molecular features^27^.

Beyond characterizing the response to a single virus, hiPSC-derived neurons allow comparison of the pathogenicity of diverse viral strains^28^. WNV strains are typically phenotyped in fibroblast- or epithelial-derived cell lines^29^, which report replication capacity but do not necessarily recapitulate infection phenotypes in neurons, the cell type most relevant for neurotropism and neuroinvasive disease. Following the introduction of WNV in North America, the ancestral NY99 genotype was rapidly displaced by the WN02 clade within a few years of introduction^30^, and WNV has continued to diversify across North America since, yet the phenotypic properties of contemporary isolates in human neurons remain largely uncharacterized.

Here we used *Ngn2*-induced WTC11 hiPSC-neurons to systematically map molecular responses to WNV infection and characterize neuron-specific replication phenotypes for contemporary WNV strains. We first established the molecular baseline of the differentiation system by temporal proteomics, then combined proteomic and transcriptomic profiling to describe the response of these neurons to WNV infection, which included cell-intrinsic immune activation, Unfolded Protein Response (UPR) induction, and declining abundance of mRNA export factors. Moreover, we found that neuronal infection by WNV is associated with calcium dysregulation and activity-dependent stress signatures, together with reduced STMN2 abundance and altered TDP-43 localization. We then used the same system, alongside non-neuronal cells such as A549, human fibroblasts, and Vero cells, to characterize three WNV isolates collected in California in 2024. Among these we identified LA24, a contemporary North American lineage 1 (WN02 clade) WNV strain with cell-type-specific replication phenotypes, including a lower replication in hiPSC-derived neurons, which was validated in primary neurons. Further supporting the neuronal phenotype observed in hiPSC, LA24 also showed reduced mortality and infection in the brain in mice.

Our study shows that this human neuronal model recapitulates the conserved antiviral and stress responses to orthoflavivirus infection while also resolving neuron-specific pathological signatures and strain-dependent replication phenotypes that are not apparent in transformed cell lines.

## Results

### Proteomic profiling defines the maturation trajectory of Ngn2-induced WTC11 hiPSC-neurons

We first sought to establish the molecular baseline of the hiPSC-derived neuron platform using proteomic profiling. Differentiation of the WTC11 hiPSC line into cortical glutamatergic-like neurons was induced by expression of murine transcription factor *Ngn2*^17,31^. To capture the transition from pluripotent state to mature-like neurons, we collected cells at days 0, 3, 7, 14, and 21 of neuronal differentiation and subjected them to global abundance proteomics quantifying over 10,000 proteins across the differentiation time course (**Fig. 1a, Supplementary Table 1**). Principal component analysis (PCA) showed a defined temporal trajectory, which accounted for 69 % of total variance in PC1, with reduced separation between day 14 and day 21 samples suggesting a plateau in remodeling of the neuronal proteome at later differentiation stages (**Fig. 1b**). Consistent with these results, the number of differentially abundant proteins relative to day 0 increased through day 14 with little additional change observed between day 14 and day 21 (**Fig. 1c**). Hierarchical clustering of proteins according to their temporal abundance profiles across differentiation stages resolved five distinct clusters, which were subjected to Gene Ontology (GO) analysis of biological processes (**Fig. 1d**). Clusters 1 (2042 proteins) and 2 (1402 proteins), containing proteins, whose abundances decreased during differentiation, were enriched in biological processes related to post-transcriptional regulation, translation and cell cycle exit. These changes are consistent with the transition from a proliferative, biosynthetically active pluripotent state to a post-mitotic neuronal identity^32^. Cluster 3 (1089 proteins) displayed a transient increase at early differentiation stages and was enriched for regulation of neuron projection development, axonogenesis, and lipid modification, suggesting an intermediate remodeling program associated with early neuronal commitment^33^. Cluster 4 (1434 proteins), containing proteins that increased during differentiation, was enriched for neuronal processes such as regulation of synapse organization or axonogenesis. As expected, cluster 4 also contained the highest proportion of brain-associated proteins, as defined by the Human Protein Atlas (**see Methods**)^34^, compared to clusters 1-3. Consistent with a trajectory from pluripotent stage to neurons, pluripotency-associated proteins, including POU5F1, NANOG, and SOX2^35^, were most abundant at day 0 and decreased rapidly following *Ngn2* induction (**Fig. 1e**). Conversely, pan-neuronal proteins such as MAP2 and TUBB3 increased during differentiation. Proteins associated with excitatory neuronal identity and synaptic maturation, including AMPA/kainate receptor subunits, glutamatergic proteins, and synaptic proteins, showed higher abundance at later time points. Metabolic proteins also increased with differentiation, consistent with the changing energetic requirements of maturing neurons^36^.

**Fig. 1.**
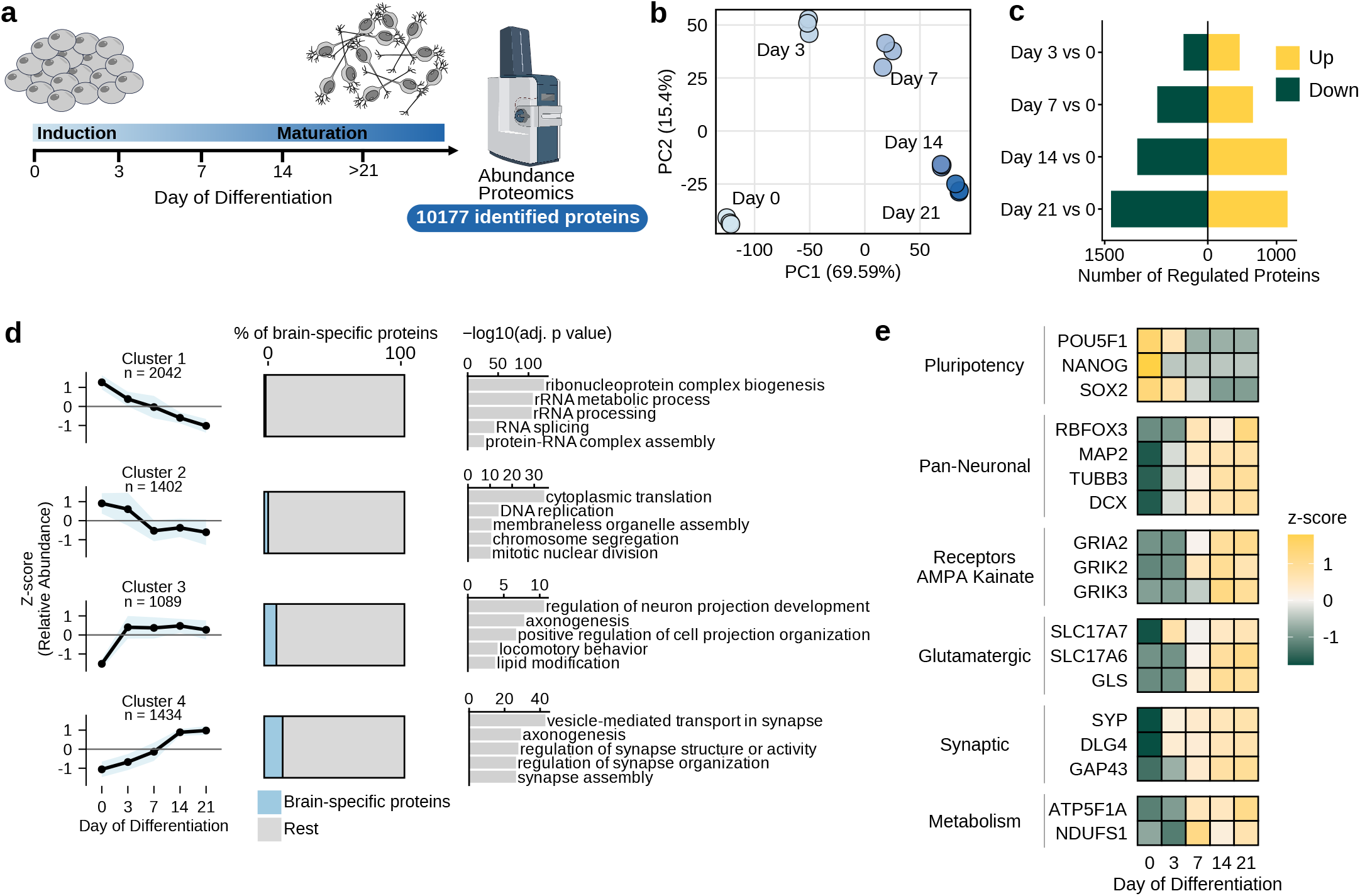
Temporal proteomic profiling of Ngn2-induced WTC11 iPSC-neurons. (a) Experimental workflow showing differentiation protocol from hiPSC to mature-like neurons. (b) Principal component analysis of proteomic profiles across differentiation. (c) Number of significantly upregulated and downregulated proteins relative to day 0. Cutoffs: log2 Fold Change: 1.5, p-value < 0.05. (d) Temporal protein-abundance clusters, proportion of brain-specific proteins, and most significantly enriched biological processes. (e) Heatmap showing the relative abundance (z-score) of selected pluripotency and neuronal protein markers. Samples from three independent differentiation replicates were collected and analyzed.

Together, these data demonstrate a rapid loss of pluripotency and cell-cycle-associated programs, followed by the acquisition of a neuronal proteomic identity and the progressive development of synaptic specialization, and establishes the proteomic baseline of the *Ngn2*-induced WTC11 hiPSC-neuron system for studies of neurotropic viral infection.

### hiPSC-derived neurons are permissive to productive WNV infection

Next, we tested if *Ngn2*-induced WTC11 hiPSC-neurons are permissive to infection with the neurovirulent WNV strain NY99 as a model^1^. Differentiated neurons at day 21 were infected with WNV NY99 at a multiplicity of infection (MOI) of 5 or 25, and infectivity and viral replication were assessed over time by performing fluorescent imaging of cells for expression of WNV NS1 protein and harvesting supernatant for plaque assays, respectively (**Fig. 2a**). Immunofluorescence analysis at 96 hours post-infection (h.p.i.) at an MOI of 25 showed widespread expression of WNV NS1 confirming that hiPSC neurons were permissive to WNV infection (**Fig. 2b**), consistent with previous results^20^.

**Fig. 2.**
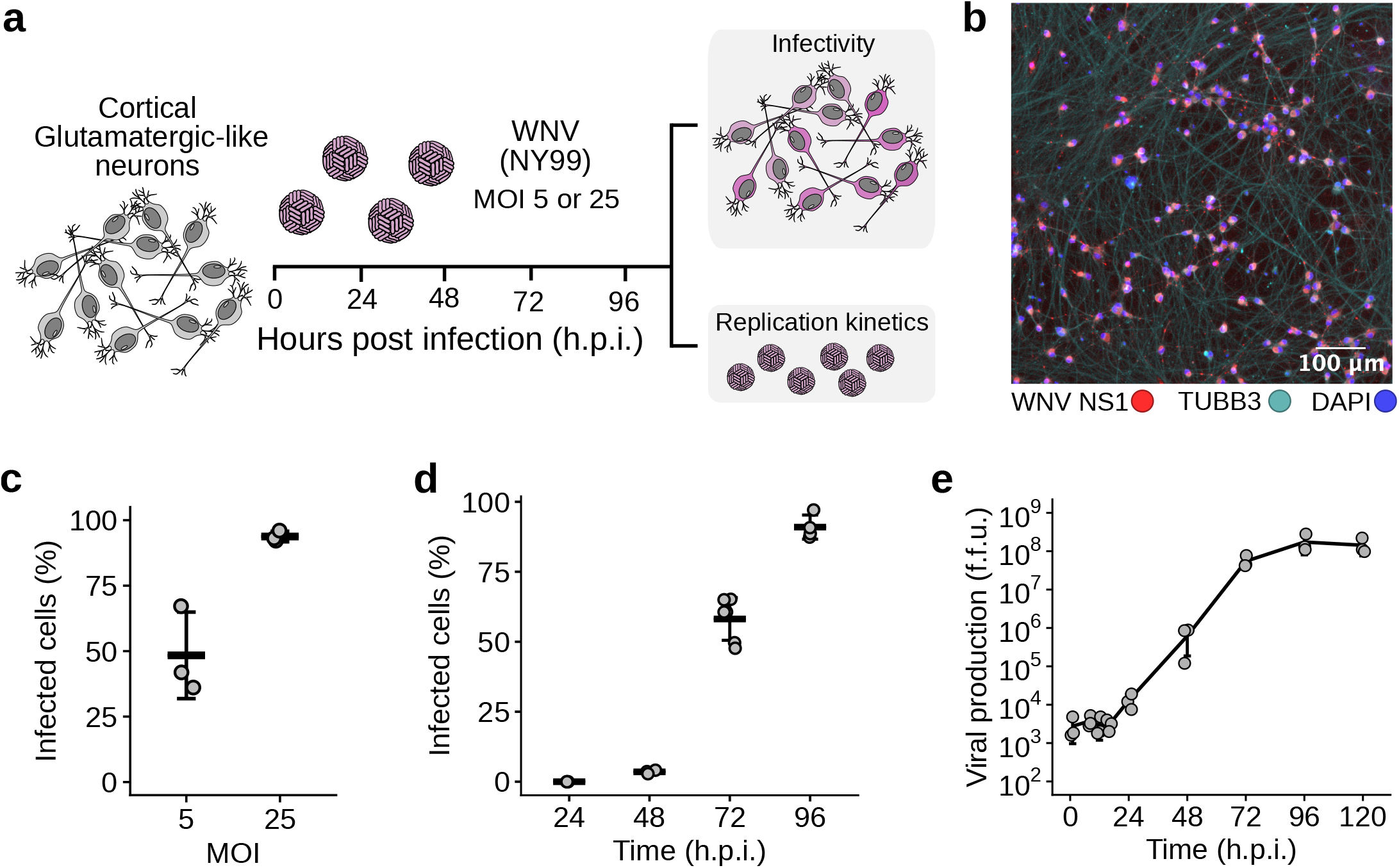
Ngn2-induced WTC11 iPSC-neurons support productive WNV infection. (a) Experimental workflow for WNV infection testing different conditions and subsequent assessment of infectivity and replication kinetics. (b) Representative immunofluorescence image at 96 h.p.i. (MOI = 25) showing WNV NS1 (red), TUBB3 (cyan), and nuclei (DAPI, blue). Scale bar, 100 μm. (c) Percentage of infected cells at 96 h.p.i. according to MOI. (d) Percentage of infected cells over time. (e) Production of infectious virus between 0 and 120 h.p.i. In (c) and (d), horizontal bars indicate the mean. In (c),(d) and (e) error bars indicate SD. In (e) the limit of detection was 10^2^ f.f.u. For (c) and (e), individual data points represent three independent differentiation replicates per condition. For (d), each data point represents the mean percentage of infected cells quantified from 3–6 images within one differentiation replicate; three independent differentiation replicates were analyzed per condition.

The proportion of infected cells increased with viral input and time. At an MOI of 5, approximately 50 % of the cells were infected at 96 h.p.i., whereas at an MOI of 25 roughly 95 % of neurons were infected at the same timepoint (**Fig. 2c**). Therefore, we selected an MOI of 25 for the following experiments. At 24 and 48 h.p.i., the proportion of infected cells remained below 10 %, before rising to approximately 60 % at 72 h.p.i. and 94.5 % at 96 h.p.i. (**Fig. 2d**). Consistent with these results, infectious-virus production increased by four orders of magnitude, reaching approximately 10^8^ focus-forming units (FFU)/mL at 96 h.p.i. achieving a plateau stage at 120 h.p.i. (**Fig. 2e**). These results demonstrate that hiPSC-derived neurons are highly susceptible to WNV infection.

### Integrated proteomic and transcriptomic profiling defines the neuronal response to WNV infection

We next combined temporal RNA sequencing (RNAseq) and abundance proteomics to characterize molecular host-cell remodeling during WNV infection. Differentiated WTC11 hiPSC-neurons (Day 21) were infected with the WNV NY99 and collected at 0, 24, 48, 72, and 96 h.p.i. Samples were subjected to bulk RNAseq and quantitative proteomics quantifying 24,688 genes and 9,377 proteins, respectively, across the time course (**Fig. 3a**)^37,38^. PCA revealed a clear time-dependent shift in both transcriptomics and proteomics datasets, with samples progressively separating in the PC1 over the course of infection (**Fig. 3b**). This separation was more pronounced at the RNA level, where PC1 accounted for 87.7 % of the total variance. In contrast, proteomic data showed a greater dispersion between replicates and smaller temporal changes over the PC1.

**Fig. 3.**
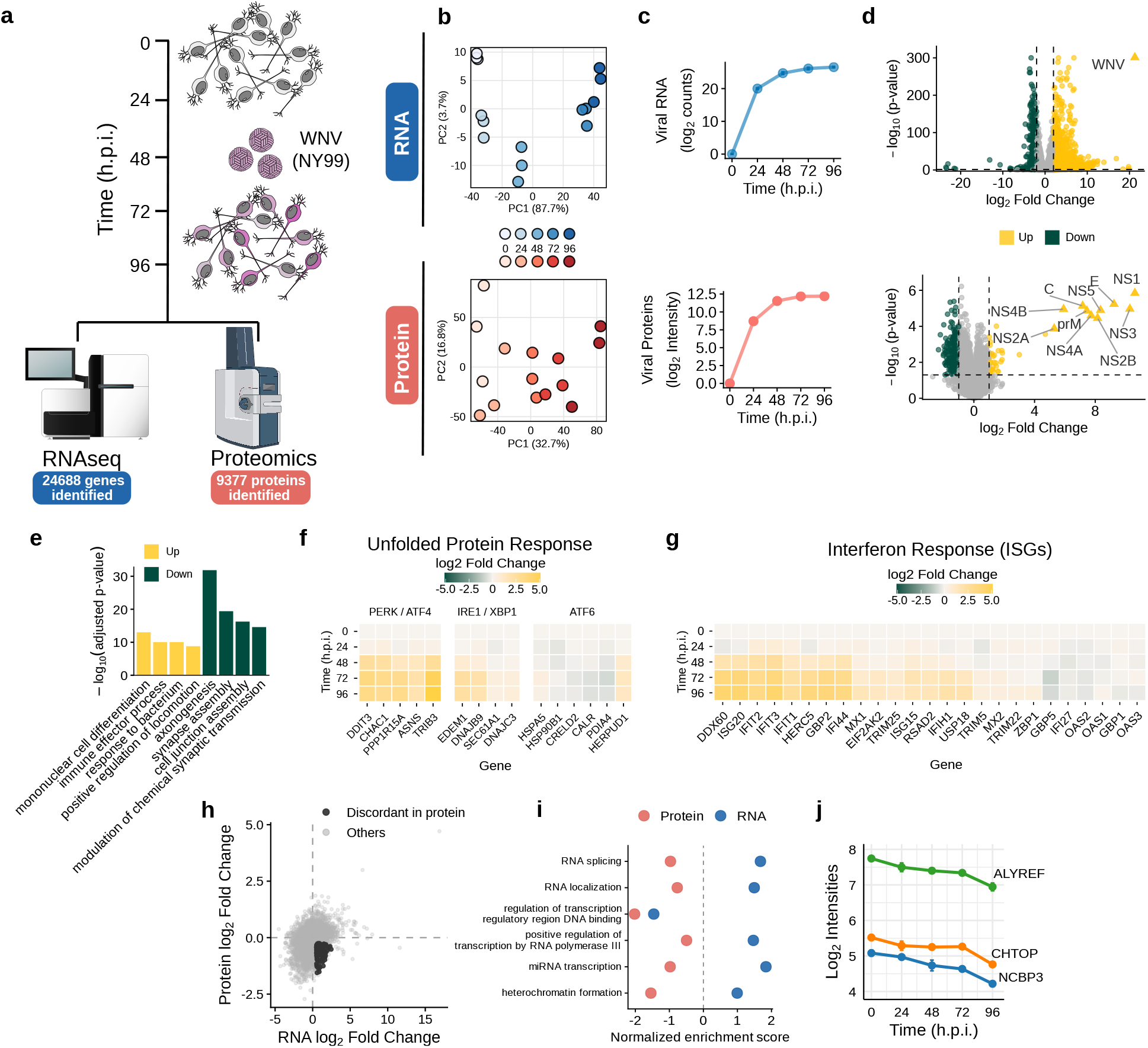
Integrated proteomic and transcriptomic profiling defines the neuronal response to WNV infection. (a) Experimental design. Differentiated WTC11 hiPSC-derived neurons were infected with WNV and collected at 0, 24, 48, 72 and 96 h.p.i. for RNA-seq and quantitative proteomic profiling. (b) Principal component analysis (PCA) of RNA-seq (top, blue) and proteomic (bottom, red) datasets. (c) Temporal accumulation of viral RNA and viral proteins during WNV infection. (d) Volcano plots showing differential RNA (top) and protein (bottom) abundance during WNV infection at 96 h.p.i. WNV genome (top) and proteins (bottom) are highlighted. Upregulated genes/proteins: yellow; Downregulated: green. (e) Functional enrichment analysis of significantly upregulated (yellow) and downregulated (green) host genes (RNA level). (f) Heatmap of transcriptomic temporal changes in representative genes associated with the three branches of the unfolded protein response. (g) Temporal expression of representative interferon-stimulated genes (ISGs) at RNA level. (h) Comparison of RNA and protein log2 fold changes. Genes displaying discordant regulation, characterized by increased transcript abundance but decreased protein abundance, are highlighted in red. Log2 Fold Change cutoffs were <-0.25 (protein) and >0.5 (RNA). (i) Gene set enrichment analysis comparing RNA- and protein-level regulation of processes enriched in the discordant set. Normalized enrichment scores (NES) are shown for the transcriptome (blue) and proteome (red). (j) Protein abundance of the mRNA export-associated factors ALYREF, CHTOP and NCBP3 across the infection time course.

Viral RNA and protein abundance increased rapidly after infection and approached a plateau between 72 and 96 h.p.i. (**Fig. 3c**), consistent with the kinetics of infection (**Fig. 2e**). Throughout the infection the individual viral proteins maintained a constant ratio across infection (**Supplementary Fig. 1a**). Differential analyses showed substantial remodeling of both the host proteome and transcriptome, together with the acute accumulation of WNV structural and non-structural proteins (**Fig. 3d, Supplementary Tables 2 and 3**). GO enrichment analysis for biological processes of regulated host genes revealed an increase in processes related to immune responses, cytokine production, and immune-cell differentiation. Conversely, downregulated genes and proteins were enriched in neuronal processes, including axonogenesis, synapse assembly, cell-projection organization, and modulation of chemical synaptic transmission (**Fig. 3e** and **Supplementary Fig. 1b**), indicating a gradual loss of neuronal processes during infection.

The transcriptional signatures above suggested that WNV triggers stress and immune programs in human neurons. We therefore focused on the gene expression of two pathways that are consistently activated by orthoflaviviruses in non-neuronal cells but whose engagement in human neurons remains poorly defined: the unfolded protein response (UPR) and type I IFN signaling. Examining markers of each UPR branch, we found the strongest induction downstream of PERK/ATF4, including *DDIT3, CHAC1, PPP1R15A, ASNS* and *TRIB3* (**Fig. 3f**). Genes associated with the IRE1/XBP1 branch, including *EDEM1* and *DNAJB9*, were also induced, whereas activation of the ATF6 branch was comparatively less and not consistent across the markers analyzed. Regarding IFN activation, WNV infection induced ISGs in hiPSC neurons in a time-dependent manner. Canonical ISGs, including *DDX60, ISG20, IFIT1, IFIT2, IFIT3, HERC5, GBP2, IFI44L, MX1, RSAD2*, and *IFIH1*, increased during infection (**Fig. 3g**)^39^. However, this response was heterogeneous, as several ISGs, including *OAS1, OAS2*, and *OAS3*, showed limited induction. This incomplete response may reflect the capacity of WNV NY99 to partially counteract antiviral responses^7,40^. Notably, *IFNA1* and *IFNA2* were undetected, whereas *IFNB1* expression was expressed at very low levels increasing modestly during infection (**Supplementary Fig. 2**).

Given that WNV infection strongly remodeled the transcriptome and proteome (**Fig. 3b**), we next asked whether changes in transcript abundance were mirrored at the protein level. Direct comparison of RNA and protein fold changes identified a subset of discordantly regulated genes whose transcripts increased while their corresponding proteins decreased (**Fig. 3h**), including the cytokine receptors IFNGR1 and IL18R1, which have been implicated in orthoflavivirus restriction (**Supplementary Fig. 3a,b**)^41,42^. Gene Set Enrichment analysis (GSEA) of discordant genes and proteins similarly revealed opposing regulation between the transcriptome and proteome for processes related to RNA splicing, RNA localization, transcriptional regulation, miRNA transcription, and heterochromatin formation (**Fig. 3i, Supplementary Fig. 2c**). These findings indicate that transcriptional induction does not necessarily translate into increased protein abundance during WNV infection, which could be a result of virus-induced translational inhibition or impaired mRNA export. Supporting the latter idea, we observed that the proteins ALYREF, CHTOP, and NCBP3, components of the TREX complex involved in mRNA export, were downregulated during WNV infection (**Fig. 3j**). Additionally, NCBP3 has been proposed to mediate selective export of specific mRNA subsets, including transcripts related to inflammatory and immune responses; its repression correlated with increased viral production for vesicular stomatitis (VSV), Semliki Forest (SFV), encephalomyocarditis (EMCV) and influenza A (IAV) viruses^43,44^. Together, our data is consistent with previous reports showing that WNV impairs nucleocytoplasmic transport^24^.

Collectively, these responses indicate that hiPSC-derived neurons are competent to viral sensing and respond to WNV infection through the canonical stress and antiviral programs described in non-neuronal cells, albeit with a partial ISG signature.

### WNV infection induces calcium dysregulation and neurodegeneration-associated stress signatures

To ask whether infection was associated with neuronal stress signatures, we tested gene sets related to neuronal excitation, excitotoxicity, and calcium response using GSEA^45^. We observed increased expression of genes related to neuronal excitation, excitotoxicity stress, and calcium-response during infection (**Fig. 4a**; **Supplementary Fig. 4**) (See methods), as exemplified by the genes *NPAS4* and *ARC*, which are tightly linked to neuronal excitation and calcium-dependent transcription (**Fig. 4b,c**)^46,47^.

**Fig. 4.**
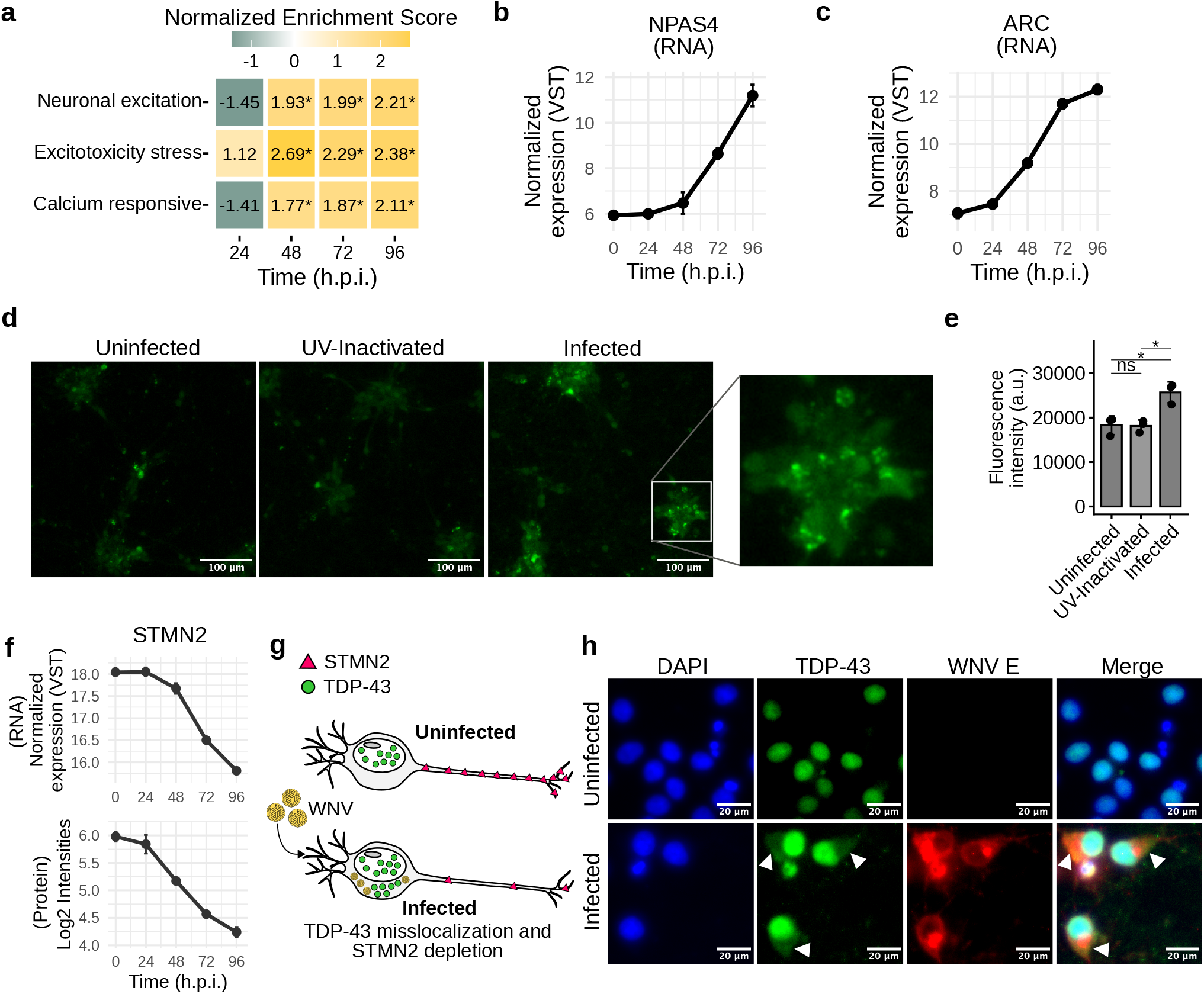
WNV infection induces calcium dysregulation and neurodegeneration-associated stress signatures. (a) GSEA showing temporal regulation of neuronal stress and calcium responsive transcriptional programs during WNV infection. Normalized enrichment scores (NES) are shown at 24, 48, 72 and 96 h.p.i. Asterisks indicate FDR < 0.05 (Benjamini-Hochberg-adjusted p-value). (b,c) Temporal RNA expression of the activity- and calcium-responsive genes *NPAS4* (b) and *ARC* (c), shown as normalized variance-stabilizing transformation (VST) expression. (d) Representative fluorescence images of intracellular calcium signal in neurons using Fluo-8 assay. Scale bars, 100 μm. (e) Quantification of intracellular calcium-sensitive fluorescence in neurons. Individual data points and mean values are shown. Individual data points represent three independent differentiation replicates per condition. ns, not significant; *P < 0.05 (two-tailed unpaired Welch’s t-tests). (f) Temporal abundance of *STMN2* RNA and protein during WNV infection. (g) Schematic representation of WNV-induced TDP-43 mislocalization and STMN2 depletion in infected neurons. (h) Representative immunofluorescence images of uninfected and WNV-infected neurons stained for DAPI (blue), TDP-43 (green) and WNV envelope protein (WNV E; red). Arrowheads indicate altered extranuclear TDP-43 localization in infected neurons. Scale bars, 20 μm. Bars in (b), (c), (e), and (f) indicate SD.

We next assessed whether the sustained neuronal excitation and ER stress during infection might lead to intracellular calcium accumulation. Using the calcium-binding dye Fluo-8, we observed that WNV-infected neurons exhibit significantly increased calcium-dependent fluorescence compared to both uninfected neurons and neurons exposed to UV-inactivated WNV (**Fig. 4d,e**). Since UV-inactivated samples did not change the fluorescence signal relative to uninfected controls, the increase in intracellular calcium was associated with productive WNV infection rather than virus exposure alone. Notably, the magnitude of the response in infected neurons was comparable to that induced by tunicamycin, an established inducer of ER stress (**Supplementary Fig. 5**), further supporting disruption of calcium and cellular stress homeostasis during WNV infection.

Because sustained Ca2+ elevation and ER stress are well-established upstream drivers of neuronal degeneration^48–50^, we evaluated whether WNV infection was accompanied by molecular features of a degenerating neuron. Stathmin-2 (STMN2), a neuron-enriched protein with important roles in axonal growth, maintenance, and repair, has been implicated in neuronal vulnerability across several neurodegenerative contexts^51^. Both STMN2 mRNA and protein abundance progressively decreased during WNV infection (**Fig. 4f**). Additionally, alterations in the location of the RNA-binding protein TDP-43, which participates in *STMN2* splicing, with its nuclear depletion and cytoplasmic redistribution, are also prominent molecular features of several neurodegenerative diseases, including amyotrophic lateral sclerosis and frontotemporal dementia^26,52^ (**Fig. 4g**). We therefore assessed whether WNV infection was accompanied by changes in TDP-43 localization. Whereas TDP-43 was predominantly nuclear in uninfected cells, infected neurons displayed altered TDP-43 distribution, including increased extranuclear/cytoplasmic signal (**Fig. 4h**). Together, these findings indicate that WNV infection induces neuronal excitation and calcium-stress responses, which coincide with reduced STMN2 abundance and altered TDP-43 localization. Thus, WNV infection-associated molecular changes overlap with features observed in neurodegenerative states.

### Isolation and phenotyping of contemporary Lineage 1 WNV strains

Having established that hiPSC-derived neurons capture infection-associated molecular changes, we next asked whether this system could also reveal phenotypic differences among naturally circulating WNV strains. Most functional characterization of WNV has relied on the NY99 reference strain, while the phenotypes of contemporary, genetically divergent isolates remain largely unexplored. To address this, we compared NY99 with contemporary WNV strains obtained from California in 2024, characterizing strain-dependent virus production across cell types in vitro, and survival and neuroinvasion in vivo (**Fig. 5a**). We isolated and sequenced three contemporary WNV isolates collected in California in 2024: LA24 (PZ920728), FRWS155 (PZ920729), and SUYA289 (PZ920730). Phylogenetic analysis placed all three isolates within the WN02 clade and among contemporary North American WNV sequences, distinct from the ancestral NY99 isolate (**Fig. 5b**).

**Fig. 5.**
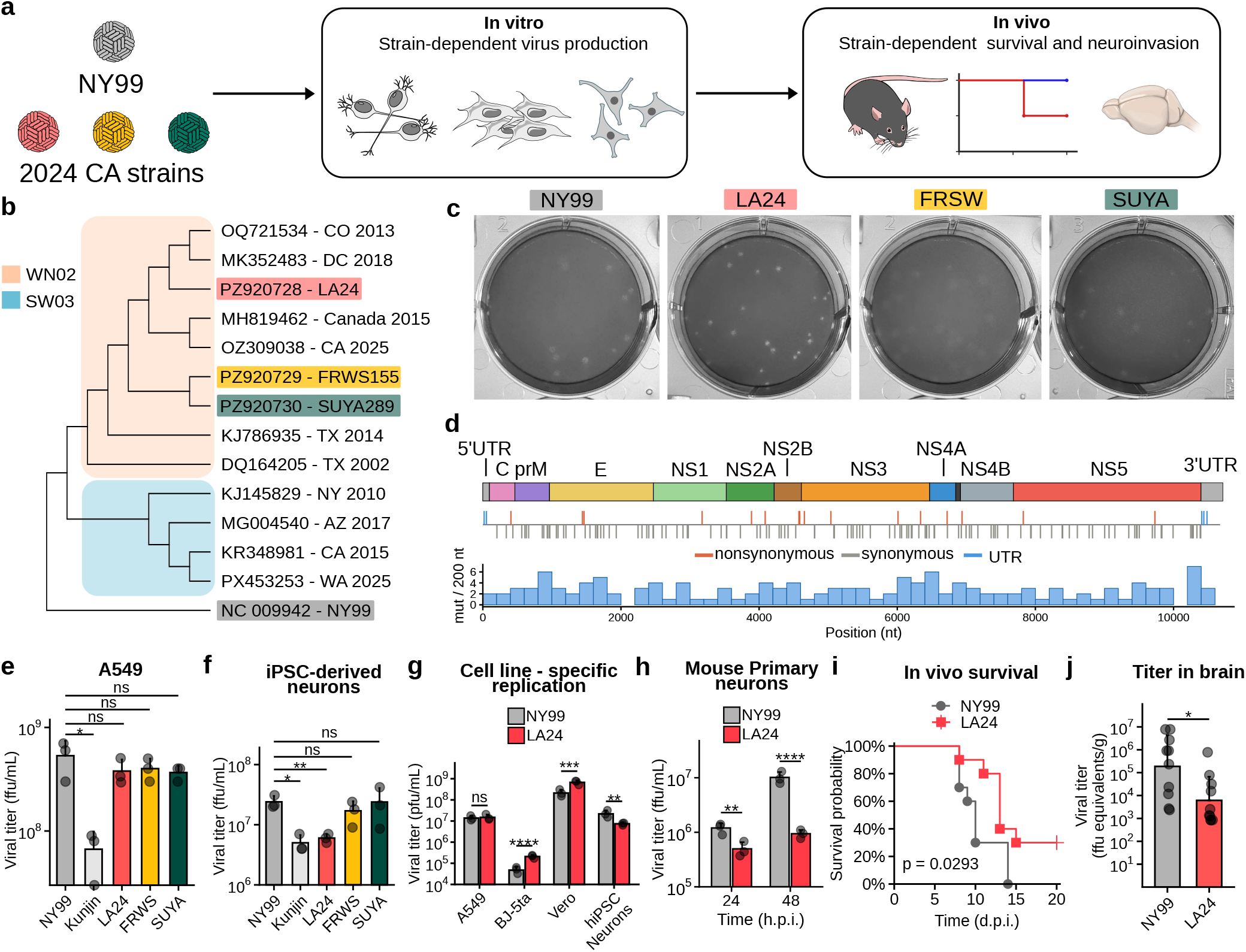
A contemporary WNV isolate exhibits neuron-specific attenuation and reduced neurovirulence in vivo. (a) Schematic of the experimental strategy used to compare the phenotypes of the ancestral NY99 strain and contemporary WNV isolates. (b) Maximum Likelihood Phylogenetic analysis of contemporary WNV isolates and other sequences of the WNV Lineage 1 group. (c) Representative plaque assays of NY99, LA24, FRWS and SUYA. (d) Genome-wide comparison of LA24 and NY99. The WNV genome organization is shown at the top, with nucleotide differences between LA24 and NY99 mapped across the genome and classified as nonsynonymous, synonymous or untranslated region (UTR) substitutions. The histogram shows the number of nucleotide differences per 200-nt window. (e) Infectious virus production in A549 cells at 48 h.p.i., MOI = 2. (f) Infectious virus production in hiPSC-derived neurons at 72 h.p.i., MOI = 25. (g) Comparison of NY99 and LA24 infectious virus production in A549 cells, BJ-5ta human fibroblasts, Vero cells and hiPSC-derived neurons. For non-neuronal cells: at 48 h.p.i., MOI = 0.1. For hiPSC-derived neurons: 72 h.p.i., MOI = 25. (h) Infectious virus production by NY99 and LA24 in mouse primary neurons at 24 and 48 h.p.i., MOI = 25. All experiments (e-h) were performed with three biological replicates. (i) Survival of mice infected with NY99 or LA24. n = 10 per group (pooled from two independent experiments). (j) Infectious viral burden in the brain of NY99- and LA24-infected mice collected at 8 days post infection (d.p.i.). n = 10 per group (pooled from two independent experiments). Symbols represent individual biological replicates where shown; bars indicate group summary values and error bars indicate standard deviation. Statistical significance is indicated in the panels; ns, not significant; *p < 0.05, **p<0.01, ***p<0.001, ****p<0.0001. For e and f, One-way ANOVA followed by Dunnett’s multiple-comparisons test was used. For g and h, Two-way ANOVA followed by Šídák’s multiple-comparisons test was used. For i, Mantel-Cox was used. For j, Mann-Whitney was used.

To determine whether these contemporary isolates displayed phenotypic variation in vitro, we performed plaque assays in Vero cells. The LA24 strain displayed reduced plaque size compared to NY99, whereas the FRWS155 and SUYA289 strains showed a more diffuse and larger plaque size (**Fig. 5c**). Focusing on LA24 we examined the extent of its genetic divergence from NY99. LA24 differed from NY99 at 143 nucleotide positions across the viral genome, corresponding to 98.7 % nucleotide identity, with synonymous and nonsynonymous substitutions distributed throughout the coding region along with changes in the untranslated regions (**Fig. 5d**). Over the ∼25 years separating the two isolates this corresponds to approximately 5.2 x 10^-4^ substitutions/site/year, in line with published estimates for WNV in North America (5.06 x 10^-4^, 95 % Highest Posterior Density (HPD) 4.44-5.70 x 10^-4^; 4.28 x 10^-4^ by root-to-tip regression)^53,54^. Applying the reported 95 % HPD interval over this period predicts approximately 122-157 substitutions, encompassing the 143 nucleotide differences observed between NY99 and LA24, which is consistent with that expected from continued WNV evolution in North America.

We next assessed whether the distinct plaque phenotype of LA24 reflected a general impairment in viral replication. In A549 cells, LA24 reached infectious titers comparable to NY99, as did FRSW and SUYA, whereas the attenuated Kunjin strain replicated to lower levels, as expected (**Fig. 5e**). However in hiPSC-derived neurons, LA24 produced significantly lower infectious titers than NY99, whereas FRSW and SUYA replicated to levels comparable to NY99 (**Fig. 5f**). To determine whether LA24 is specifically attenuated in neurons, we compared NY99 to LA24 viral production in an expanded set of non-neuronal cell types including human fibroblasts (BJ-5ta) and Vero cells. LA24 produced similar (A549) or significantly higher amounts of infectious particles (BJ-5ta and Vero) in non-neuronal cells, whereas infectious titers were consistently lower in hiPSC-neurons compared to NY99 (**Fig. 5g**). Furthermore, LA24 produced significantly lower viral titers than NY99 in primary mouse neurons (**Fig. 5h**), demonstrating that the attenuated replication phenotype was conserved across independent human and murine neuronal models.

Finally, we asked whether the neuronal phenotype identified in vitro could be translated to differences in pathogenesis in vivo. Mice infected with LA24 showed significantly increased survival compared with NY99-infected animals (p=0.0293; **Fig. 5i**). Consistent with this reduced disease severity, infectious viral burden in the brain was significantly lower for LA24 than NY99 at 8 days post infection (**Fig. 5j**). Together, these findings identify LA24 as a naturally circulating WNV isolate with an attenuated phenotype in neurons, and associated with reduced pathogenicity in vivo.

More broadly, these results highlight the capacity of hiPSC-derived neurons to serve as a platform for defining the neuron-specific molecular responses to viral infection and identifying naturally occurring viral variants with cell type-specific phenotypes.

## Discussion

Neuronal injury is a defining feature of WNV disease; however, the molecular responses to WNV infection in human neurons remain poorly defined. In this study, we use hiPSC-derived neurons (**Fig. 1**) as a scalable system to examine the cell-intrinsic consequences of neurotropic viral infection in a disease-relevant cell type (**Fig. 2**). Using this platform, we first defined the molecular responses of human neurons to WNV infection through temporal RNAseq and proteomics (**Fig. 3**). With 9,377 protein groups quantified, this is, to the best of our knowledge, the first temporal proteome of WNV infection in human neurons and the deepest reported for WNV^37,38^. This analysis revealed a partial type I IFN signature, a UPR dominated by PERK/ATF4, uncoupling of transcript and protein abundance coinciding with depletion of TREX mRNA export factors, and progressive loss of synaptic and axonogenesis proteins alongside calcium dysregulation (**Fig. 4**). Applying the same system to contemporary isolates, we identified LA24, a WN02-clade lineage 1 strain whose attenuated replication was apparent in hiPSC-derived neurons but not in A549, fibroblast or Vero cells (**Fig. 5**). This neuron-restricted phenotype was reproduced in primary mouse neurons and tracked with increased survival and lower brain viral burden in vivo. Together, these findings establish hiPSC-derived neurons as a platform for both mechanistic dissection of neuronal responses to WNV and functional discrimination of viral strains, including phenotypic differences that are not readily captured in transformed cell lines.

Our first goal was to define the proteomic features of *Ngn2*-driven differentiation in the WTC11 model, and establish a baseline of cellular responses against which infection-induced changes could be distinguished. Temporal proteomic profiling of NGN2-driven differentiation has been reported from hESCs^55^, from iPSC-derived glutamatergic and motor neurons^56^, and for the KOLF2.1J line together with its phosphoproteome^57^. In the WTC11 cell background, which underlies the widely used i^3^Neuron platform, only a two-state comparison between iPSCs and differentiated neurons has been performed so far^58^. Our dataset defines the differentiation trajectory in this experimental background, quantifying 9,126 proteins across five time points and capturing the transition from a pluripotent state to a post-mitotic neuronal identity, with a transient increase in proteins linked to early neuronal commitment, and the progressive acquisition of synaptic and metabolic maturation markers.

Against this baseline, we obtained a temporally resolved proteomics and transcriptomics map of WNV infection, which revealed two coordinated signatures: induction of immune and cytokine programs, and progressive loss of proteins governing axonogenesis, synapse assembly and synaptic transmission. A comparable inhibition of synaptic pathways has been reported in mouse neurons infected with WNV and JEV^59^, suggesting that loss of neuronal function precedes cell death. The IFN response was measurable but selective: canonical ISGs were induced, whereas the OAS cluster was not. IFN-alpha transcripts were undetected and *IFNB1* was expressed at very low levels, increasing only modestly during infection, consistent with reports that human and murine neurons are poor producers of type I IFN even when they respond well to it^60,61^. This suggests that much of the ISG induction we observe may arise from direct, IRF3-dependent transcription downstream of RIG-I/MDA5 sensing of WNV RNA, which has been shown to restrict WNV through both IFN-dependent and IFN-independent mechanisms^62,63^. Since bulk measurements cannot distinguish whether ISGs are induced in infected neurons or in neighbouring uninfected cells, future work will address whether the neuronal injury we observe is restricted to virus-infected neurons or extends to uninfected bystander cells.

An additional pattern during WNV infection was the uncoupling of transcript and protein abundances for a set of genes, as revealed by GSEA for modules associated with transcriptional regulation. TREX-associated factors such as NCBP3, ALYREF and CHTOP decreased in abundance at protein level over the infection course. Because NCBP3 can exert antiviral effects by facilitating the export of antiviral transcripts^43,44^, disruption of nucleocytoplasmic transport might provide a selective advantage for orthoflavivirus replication. This idea is consistent with previous studies showing that orthoflaviviruses interfere with host mRNA processing, including the degradation of nucleoporins by the DENV and ZIKV NS2B-NS3 proteases, which compromises nuclear export^24^. Infection also activated the UPR; the response was dominated by the PERK/ATF4 branch, with moderate IRE1/XBP1 and limited ATF6 activation, which contrasts with the attenuated Kunjin strain, which elicits a strong ATF6 branch activation, although we only assessed UPR at the abundance level^23,64^. Since PERK activation results in translation inhibition through eIF2α phosphorylation, this offers another potential explanation for the protein-transcript discordance described above. More studies focused on transcript localization and translational output will be needed to resolve the contribution of impaired mRNA export and UPR-mediated translational arrest.

The upregulation of genes responsive to calcium levels pointed to altered excitatory signaling and a potential disruption of calcium homeostasis, accompanied by induction of the activity-dependent genes *NPAS4* and *ARC*. We confirmed that intracellular calcium rose to levels comparable to tunicamycin treatment and required productive WNV infection, since UV-inactivated virus had no effect. The increase in intracellular calcium levels is consistent with calcium release from the ER downstream of the UPR activation, altered plasma-membrane permeability and excitotoxic signalling. Within the stress context, STMN2 declined at both mRNA and protein level, and TDP-43 showed increased extranuclear signal. Although the connection could be causal, STMN2 levels can also be rapidly depleted during acute stress, independently of TDP-43 splicing^65^. TDP-43 redistribution is nonetheless a recurring outcome of neurotropic infection, reported for enteroviruses D68 and A71, and Theiler’s murine encephalomyelitis virus^66–68^, and is of interest given epidemiological associations between viral encephalitis and later neurodegenerative diagnosis^27^.

Our data are relevant to questions about the intrinsic susceptibility of human neurons to WNV. A recent study in neural cultures derived from human fetal neural progenitors reported that neurons were largely refractory to infection, with productive replication confined to astrocytes and oligodendrocytes and neuronal death occurring through bystander mechanisms rather than direct infection^28^; notably, JAK/STAT inhibition dramatically increased infection in glial cells without restoring neuronal infection, arguing against an IFN-mediated block as the basis for neuronal resistance. In contrast, we found that hiPSC-derived cortical glutamatergic neurons support productive replication and mount a coordinated cell-intrinsic response to infection, a conclusion reinforced by the requirement for replication-competent virus in our calcium experiments. Our observations are consistent with reports of productive WNV infection in iPSC-derived cortical and spinal motor neurons^20^, and with the localization of viral foci to neuron- and astrocyte-rich regions of human cerebral organoids^14,15,69^. Several non-exclusive factors may reconcile these findings, including neuronal sub-type, differentiation protocol and maturation state, the presence or absence of glia, and the MOI and duration of infection. Defining the cell-intrinsic determinants of neuronal permissiveness, and how they vary across neuronal subtypes, remains an open question that genetically tractable hiPSC systems are well placed to address.

Finally, we used this platform to identify that a contemporary lineage 1 strain, LA24, showed neuron-specific attenuation. The attenuation of LA24 in our neuronal model, recapitulated in mouse primary neurons, was also associated with increased survival and decreased viral burden in the brain in mouse infection studies. LA24 differs from NY99 at 143 nucleotide positions, but which of these differences underlie the cell-type-specific phenotype remains to be elucidated. Resolving these functionally relevant differences in sequences could help define the viral features contributing to neuronal tropism.

Our study defined the trajectory of neuronal response to WNV infection and identified a naturally attenuated contemporary strain; however, several limitations should be considered. First, our system consists of neurons in monoculture, without the presence of other glial cells such as astrocytes or microglia, which can affect neuron maturation; consequently, we did not capture immune cell responses to infection or other glial contributions. Second, the study was performed on one exclusive genetic background using a defined MOI and time-points of infection, and using cortical glutamatergic neurons. Future work could extend this platform to other MOIs and relevant neuronal cell types. Despite these limitations, our study provides a robust framework for WNV infection in human neurons, as well as a new strain with cell-type specific tropism.

## Methods

### Cells

Vero E6 (African green monkey kidney, ATCC CRL-1586), A549 (human lung carcinoma, ATCC CCL-185) and BJ-5ta (ATCC CRL-4001) cells were obtained from the American Type Culture Collection (ATCC). Cells were maintained in Dulbecco’s Modified Eagle Medium (DMEM, Gibco) supplemented with 10 % heat-inactivated fetal bovine serum (FBS, Thermo Fisher). Cultures were incubated at 37 °C and 5 % CO2.

### hiPSC culture

Doxycycline-inducible murine *Ngn2* WTC11 hiPSCs were obtained from the Li Gan Lab^17^. iPSCs were grown on Matrigel-coated plates in Essential 8 Flex medium (Gibco). Accutase (BioLegend) was used for dissociation. Chroman I (MedChemExpress) was used as a Rock inhibitor for replating.

### hiPSC differentiation into neurons

For induction (D0-D3), DMEM/F12 (Gibco) supplemented with doxycycline (2 μg/mL, Fisher Scientific), 1X N2 supplement (Thermo Fisher), 1X Glutamax (Gibco), 1X NEAAs (Gibco), and 50 nM Chroman I (only on D0) was used. On D3, predifferentiated neurons were dissociated using Accutase and replated at 5e6 cells per well in MW6 plates coated with 20 μg/mL of poly-D-lysine (Thermo Fisher) and 2 μg/mL of laminin (Fisher Scientific), using D3 maturation medium: DMEM/F12:BrainPhys (StemCell Technologies) (1:1), Chroman I (50 nM), 1X N21-max supplement (RandD), 1X BDNF (10 ng/mL, Thermo Fisher), 1X GDNF (10 ng/mL, Thermo Fisher), NT3 (10 ng/mL, Thermo Fisher), doxycycline (2 μg/mL). On D4, the medium was completely replaced with D4 maturation medium until D7, consisting of the D3 formulation omitting doxycycline and Chroman I, and supplemented with 2 μM Ara-C (Sigma). At D7, the medium was replaced with Ara-C-free D4 maturation medium until D10. From D10 to D21, half media changes every 3 to 4 days, using BrainPhys media supplemented with 1X N21-max supplement, 1X BDNF (10 ng/mL), 1X GDNF (10 ng/mL), NT3 (10 ng/mL).

### Mouse primary neurons

Primary cortical neurons were prepared from P0 mouse cortices. Cortices were dissected in ice-cold HBSS and digested with papain (10 U/mL, Worthington) for 25 min at 37 °C. Following enzymatic digestion, the tissue was gently triturated and passed through a 70 µm cell strainer. Dissociated neurons were plated onto Matrigel-coated coverslips in 24-well plates in MEM-based plating medium (Gibco) containing 0.5 % glucose (MilliporeSigma), 0.02 % NaHCO_3_ (Gibco), 0.1 mg/mL transferrin (MilliporeSigma), 10 % FBS (Thermo Fisher), 2 mM L-glutamine (Gibco), and 0.025 mg/mL insulin (MilliporeSigma). On day in vitro (DIV)1, approximately 90 % of the plating medium was replaced with Neurobasal-based growth medium (Gibco) containing 0.5 % glucose, 0.02 % NaHCO_3_, 0.1 mg/mL transferrin, 5 % FBS, 2 % B27 supplement (Gibco), and 0.5 mM L-glutamine. Cultures were infected with WNV on DIV11 with a MOI of 25.

### Viruses

WNV strain NY99 and Kunjin were obtained from BEI resources (Genbank AY842931 and KX394396 respectively).

For the isolation of WNV strains (PZ920728, PZ920729 and PZ920730), mosquito pools collected in California in 2024 were homogenized in 1 mL of medium. The homogenate was clarified by centrifugation at 10,000 x g for 10 min at 4 °C. The resulting supernatant was filtered (0.22 µm) and inoculated onto subconfluent Vero cell cultures for a 1 h adsorption period. Cultures were monitored daily for cytopathic effect (CPE) for up to 5 days. Supernatant was collected as passage 1.

WNV stocks were propagated as previously described^70^. Viral titers were determined by standard plaque assay and fluorescence plaque assay in Vero, using anti-envelope (E) D1-4G2-4-15 antibody (Sigma-Aldrich). Only passage 1 and 2 viral stocks were used in this work.

WNV infections in hiPSC-derived neurons were performed using a MOI of 25 unless otherwise specified.

### Animal ethics statement

All animal experiments and procedures were performed in accordance with the recommendations in the Guide for the Care and Use of Laboratory Animals of the National Institutes of Health, Bethesda, MD. The protocols were approved by the Institutional Animal Care and Use Committee at the Washington University School of Medicine (assurance number A3381-01). Injections were performed under anesthesia that was induced and maintained with ketamine hydrochloride and xylazine, and all efforts were made to minimize animal suffering.

### Survival analysis

Four-week-old male C57BL/6J mice (Jackson Laboratories, 000664) were inoculated by intraperitoneal injection with 100 FFU of WNV-NY99 or WNV-LA24 and monitored daily for survival. Survival distributions were compared using the Kaplan–Meier method and the Mantel–Cox log-rank test. 10 mice for each group were used by pooling data from two independent experiments.

### Viral burden analysis

Four-week-old male C57BL/6J mice were inoculated by intraperitoneal injection with 100 FFU of WNV-NY99 or WNV-LA24, and studies were terminated at day 8 after infection. Mouse brains were harvested after perfusion with 25 mL of dPBS and titered by qRT-PCR using RNA isolated from viral stocks as a standard curve to determine focus-forming unit (FFU) equivalents. RNA extraction was performed using the KingFisher Flex system (Thermo Fisher Scientific). WNV RNA levels were quantified by one-step quantitative reverse-transcription PCR (RT-qPCR) using a TaqMan probe-based assay targeting the WNV envelope (E) gene. The assay utilized the Taqman RNA-to-Ct 1-step kit (Thermo Fisher Scientific) along with an untranslated region and NS1 specific primer/probe set^71^, using the following oligonucleotides:

~~~
F:5’-TCAGCGATCTCTCCACCAAAG-3’
R:5’-GGGTCAGCACGTTTGTCATTG-3’
Probe:5’-/56-FAM/TGCCCGACCATGGGAGAAGCTC-TAMRA-3’
~~~

WNV RNA abundance was quantified relative to a standard curve generated using serial dilutions of a WNV RNA standard and expressed as viral RNA copies per gram of tissue. Appropriate negative extraction controls and no-template controls were included in each experiment. Viral titers in brain tissue were compared between groups using the two-tailed Mann–Whitney U test.

### Immunofluorescence assays

For IFA, neurons were fixed in 4 % PFA (10 min at RT, Sigma) and rinsed with 1X DPBS (Fisher Scientific). Cells were blocked and permeabilized using 20 % FBS and 0.1 % Triton X-100 in DPBS. Primary antibodies were incubated 1 h at 4 °C, followed by secondary antibodies for 30 min at RT. Nuclei were strained with DAPI (1:3000) (D1306, ThermoScientific) for 30 min. Images were captured using a Discover Echo Revolve microscope and processed using ImageJ version 1.54p.

Rabbit polyclonal anti-β-III-tubulin (PRB-435P, BioLegend) was used at 1:1000. Mouse monoclonal anti-WNV NS1 antibody (used at 1:500) was obtained through BEI Resources, NIAID, NIH: Monoclonal Anti-West Nile Virus Nonstructural Protein 1, Clone 22-NS1 (produced in vitro), NR-10145^72^. Rabbit polyclonal anti-TDP-43 antibody was used at 1:1000 (10782-2-AP, Proteintech). For secondary antibodies, antimouse Alexa Fluor 546 (A11030, Invitrogen), anti-rabbit Alexa Fluor 488 (A32731, Invitrogen), anti-rabbit Alexa Fluor 647 (A32795, Invitrogen) were used at 1:2000.

### Calcium assay

Calcium Flux Assay Kit (Fluo-8, No Wash) (ab112129, Abcam) was used as specified by the manufacturer, with an incubation of 30 min at RT, and fluorescence was measured using a CLARIOstar (BMG Labtech) plate reader. hiPSC-derived neurons were differentiated in 96 well plates for 21 days and either remained uninfected, or were infected with WNV for 72 h. Tunicamycin was added at 1 µg/mL 24 h before the assay (T7765, Sigma-Aldrich). DMSO was used as a vehicle for the uninfected control. UV-inactivated virus, generated by irradiating viral stock with UV light for 3 h and confirmed to be non-infectious by testing in Vero cells, was included as an additional control.

### Sequencing of viral isolates and genome assembly

RNA from infected cells was extracted using Quick-RNA Miniprep Plus Kit (Zymo Research). RNA concentration and quality was measured using Agilent TapeStation (RNA HS ScreenTape) ensuring high-quality input (RIN ≥ 8.8). Sequencing libraries were generated from 100 ng of total RNA using the NEBNext Ultra II Directional RNA Library Prep Kit, supplemented with ERCC Spike-in (External RNA Controls Consortium). RNA was fragmented for 8 min at 94 °C to target appropriate insert sizes. Following first and second-strand synthesis, cDNA underwent a 1.8x SPRI bead cleanup (AMPure XP). Libraries were then end-repaired, ligated to NEBNext adapters (1:25 dilution), and purified with 0.9x SPRI beads (Beckman Coulter). Final library amplification was performed using 9 cycles of PCR with barcoded primers and USER enzyme digestion. The resulting libraries were purified with 0.8x SPRI beads and quantified for downstream sequencing. Barcoded libraries were pooled and sequenced on an Illumina NextSeq 2000. Raw sequencing reads underwent quality assessment via FastQC, followed by adapter removal and quality filtering using Trimmomatic v0.39 (parameters: ILLUMINACLIP:TruSeq3-PE:2:30:10, SLIDINGWINDOW:4:15, and MINLEN:36).

### RNAseq of WNV-infected neurons

For library preparation, 50 ng of purified RNA from each sample was used as input. An ERCC spike-in mix (External RNA Controls Consortium) was added to each sample as an internal positive control. To deplete ribosomal RNA from mammalian transcripts, FastSelect-rRNA HMR (Qiagen) was included at a 1:10 dilution. Libraries were prepared using the NEBNext Ultra II Directional RNA Library Prep Kit (New England Biolabs, E7760S), following the manufacturer’s protocol on the firefly liquid handler platform (sptlabtech). Briefly, RNA was reverse-transcribed into cDNA, which was then used for construction and barcoding of sequencing libraries. Final libraries were sequenced using 146-nucleotide paired-end reads on the Illumina NovaSeqX 10B, with a target sequencing depth of at least 50 million reads per sample.

Raw sequencing reads were assessed for quality using FastQC (v0.12.1) and aggregated with MultiQC (v1.32)^73^. Adapter and quality trimming was performed with fastp (v1.0.1) using default parameters^74^. Transcript-level quantification was carried out with Salmon (v1.10.3)^75^, using selective alignment^76^, with sequence-specific (--seqBias) and GC (--gcBias) bias correction enabled and automatic library-type detection (-l A; inferred as ISR). Indices were built with k = 31 from a FASTA combining the human reference transcriptome (GENCODE v44, GRCh38, Ensembl 110)^77^, with the genome sequence of WNV NY99 (NC_009942.1); samples were mapped against the index corresponding to WNV NY99 genome. Transcript-level quantifications were imported into R (v4.5.2; Bioconductor v3.22) with tximport (v1.38.2)^78^, using countsFromAbundance = “lengthScaledTPM”. Differential expression analysis was performed with DESeq2 (v1.50.2)^79^. Benjamini–Hochberg correction was applied for the contrasts (p-adjusted value < 0.05).

Genes selected for GSEA were manually curated based on bibliography (Supplementary Table 4)^80,81,82,46,47,83,84,85,86,87,88,89,90,91, 92,93,94,95,96,97,98,99,100,25,101,102^. GSEA was performed using the fgsea package (v1.36.2)^103^. Genes were ranked according to the Wald statistic obtained from DESeq2 for each condition relative to the 0 h reference. Gene sets related to neuronal excitation, calcium-responsive signaling, and excitotoxicity/stress responses were tested using the fgsea-Multilevel algorithm^104^. Normalized enrichment scores (NES) and Benjamini–Hochberg-adjusted p values were used to assess pathway enrichment.

### Proteomics sample preparation and LC-MS

hiPSC-derived neurons were collected in triplicates. For proteomics sample preparation, we used the previously described SPEED protocol^105^. Briefly, cell pellets were lysed using 100 % trifluoro acetic acid (TFA, Fisher Scientific) and neutralized with 2 M TrisBase. Samples were reduced by incubation for 5 min at 95 °C with Tris(2-carboxyethyl)phosphine (TCEP, Sigma) and 2-Chloroacetamide (CAA, Sigma) (100 mM and 400 mM respectively). Lysates were digested with trypsin (Promega) and lysC (FUJIFILM Wako Chemicals) for 20 h at 37 °C at 1:50 (enzyme:protein w:w). Next, C18 cleanup was performed: columns (70 μg, Higgins) were activated with 200 μL methanol (Fisher Scientific), washed with 200 μL (x3) 80 % acetonitrile (ACN, Fisher Scientific) / 0.1 % Trifluoroacetic Acid (TFA), equilibrated with 200 μL (x3) of 2 % ACN / 0.1 % TFA. Columns were washed with 200 μL (x3) of 2 % ACN / 0.1 % TFA, and samples were eluted with 75 μL (x2) 50 % ACN/ 0.1 % TFA. Samples were dried in a vacuum centrifuge and resuspended in 0.1 % formic acid (FA).

Samples were analyzed on a timsTOF HT mass spectrometer (Bruker) coupled to a Vanquish Neo UHPLC system (Thermo Fisher Scientific). Peptides were separated at 500 nL/min using a nonlinear gradient from 2 to 80 % of 0.1 % FA / 80 % ACN buffer over 30 min. Data was acquired in dia-PASEF mode using a 75 ms TIMS ramp and a 75 ms Accumulation time. The TIMS precursor MS window covered an *m/z* range of 100-1700 Da and a 1/K_0_ range of 0.72-1.50. One TIMS precursor MS frame was followed by 10 dia-PASEF MS2 frames using fragmentation windows having a width of 30 Da with an m/z overlap of 1 Da between windows. Fragmentation windows spanned a m/z range of 250-1325 Da and a 1/K_0_ range of 0.68-1.40.

### Proteomics data analysis

Mass spectrometry data were analyzed with Spectronaut (v20.5, Bio-gnosys) using directDIA. Spectra were searched against the UniProt human canonical proteome (20,405 entries, downloaded 2 July 2025), the WNV NY99 proteome (10 entries), and a universal contaminant database (248 entries). Peptide fragment ion intensities were exported from Spectronaut in .tsv format and imported into R (v4.5.2) for downstream analysis using the MSstats framework (v4.18.1)^106^. Contaminant entries were removed, and reports were converted to MSstats format using SpectronauttoMSstatsFormat with a precursor q-value cutoff of 0.01, retaining proteins identified by a single feature and including non-unique peptides so that shared viral peptides were preserved. For viral proteins, zero intensities were converted to missing values prior to summarization to allow them to be treated as censored. Feature-level intensities were log2-transformed, normalized by median equalization, and summarized to the protein level with dataProcess using Tukey’s median polish, with all features retained (featureSubset = “all”, min_feature_count = 1); missing values were treated as censored (censoredInt = “NA”) and imputed with the accelerated failure time model (MBimpute = TRUE, maxQuantileforCensored = 0.999). UniProt accessions were mapped to gene symbols using UniProt.ws; viral proteins were annotated with their protein name.

Brain proteins were defined on proteins expressed exclusively in brain tissue by mass spectrometry in the Human Protein Atlas (HPA, v25.1)^34^.

For differential abundance, protein log2 intensities were analysed with limma (v3.66.0)^107^. A design matrix was constructed without an intercept on the differentiation timepoint factor (D0, D3, D7, D14, D21), and linear models were fitted with lmFit. Contrasts of each time-point against D0 were defined with makeContrasts and t-statistics were computed with eBayes. P-values were adjusted using the Benjamini– Hochberg procedure. Proteins with P < 0.05 and |log2FC| > 1 in at least one contrast were considered significant and retained for clustering. For clustering, log2 intensities were averaged across replicates within each timepoint and each protein profile was standardized to a z-score across the five timepoints. Clustering was performed with Mfuzz (v.2.70.0)^108^.

### Statistical analysis

Viral titers in brain tissue were compared between groups using the two-tailed Mann–Whitney U test. Statistical analyses for mice experiments were performed using GraphPad Prism, and P < 0.05 was considered statistically significant.

For Fluo-8 calcium assay, comparisons between conditions (3 biological replicates) were made using two-tailed unpaired Welch’s t-tests.

## Data availability

Processed data supporting the findings of this study are provided in the Supplementary Tables accompanying this manuscript. Raw proteomics and transcriptomics data generated in this study will be deposited in appropriate public repositories. Repository accession numbers will be provided in a revised version of the preprint and in the final published article.

## Acknowledgments

We thank Jan Carette for many helpful discussions.

## Funding

A.M. was supported by the Chan Zuckerberg Biohub San Francisco Postdoctoral Fellowship and by the Stanford Synthetic Biology Program.

## Author contributions

A.M., R.H. and C.A. conceived and designed the study. A.M. performed the majority of the experiments, with contributions from A. Anaya, A. Akgul, N.W., A.S., S.L. and J.W. A.M. analyzed the data and prepared the figures, with input from C.A. and R.H. M.W., T.C.S., C.M.B., M.S.D., C.A. and R.H. supervised the work. A.M. and R.H. acquired funding. A.M., C.A., and R.H. wrote the manuscript. All authors read and corrected the article and contributed to the interpretation of results.

## Competing interests

M.S.D. is a consultant or advisor for Inbios, IntegerBio, Akagera Medicines, Moderna, Merck, GlaxoSmithKline, and Bird & Bird LLP. The Diamond laboratory has received unrelated funding support in sponsored research agreements from Moderna.

---

**Supplementary Fig. 1.**
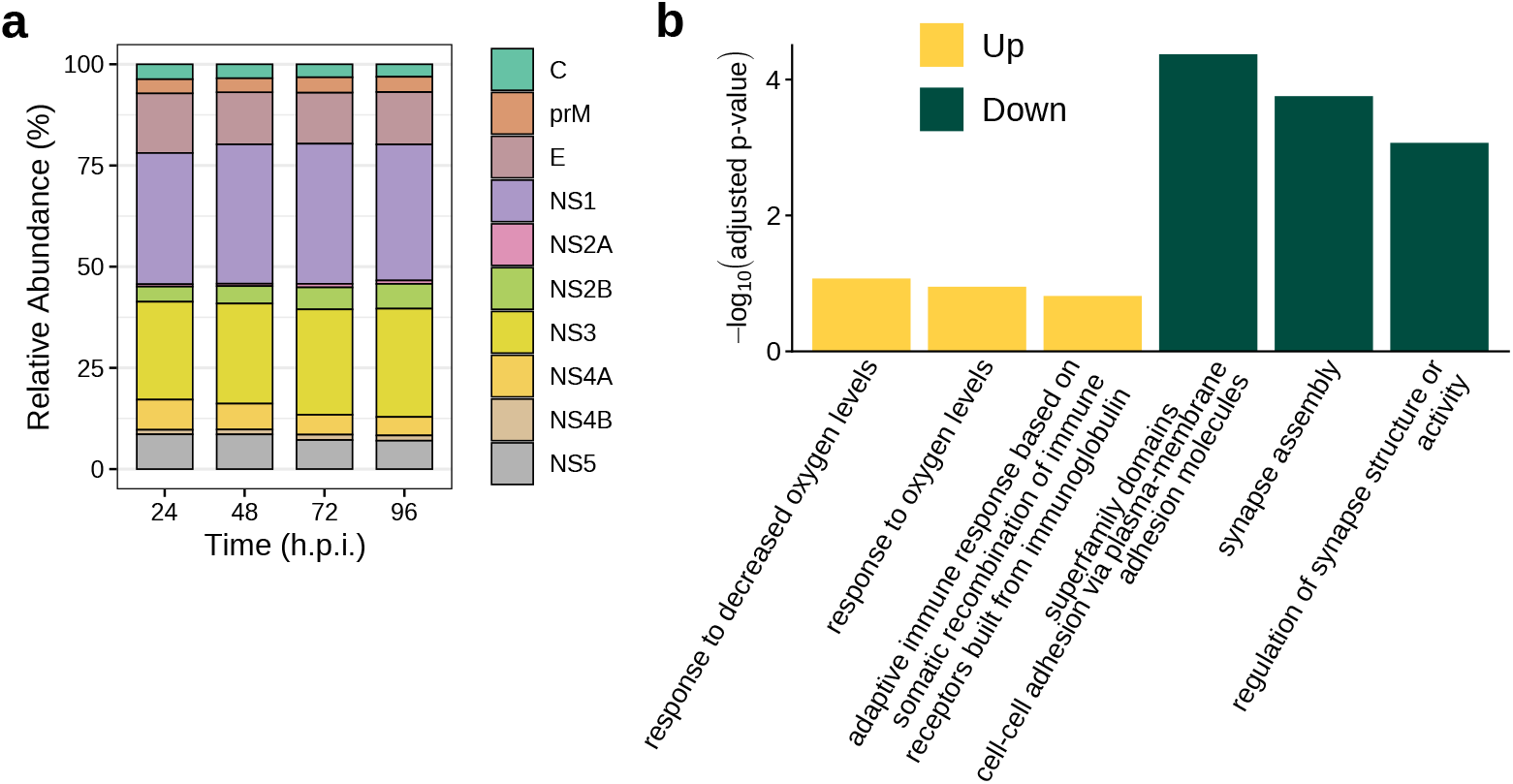
Proteomics of neuronal response to infection. (a) Relative abundance of individual WNV proteins at 24–96 h.p.i. (b) Gene Ontology enrichment analysis of upregulated or downregulated proteins, showing representative significantly enriched biological processes.

**Supplementary Fig. 2.**
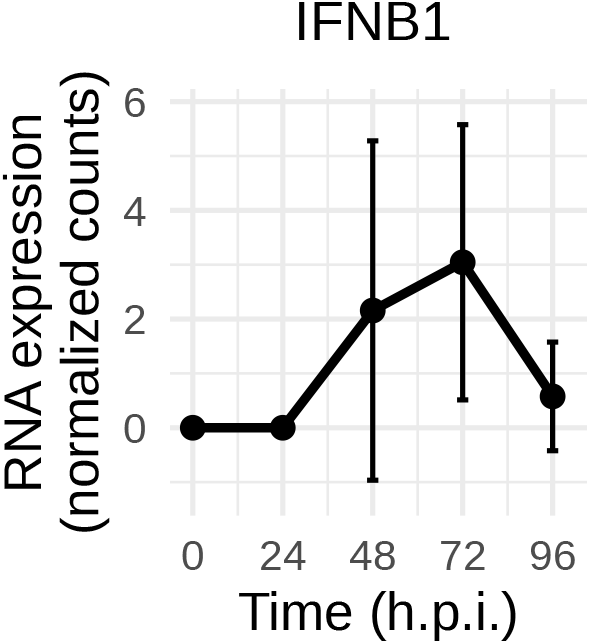
Type I IFNB1 expression in WNV-infected hiPSC-derived neurons. RNA-seq normalized counts for *IFNB1* expression in hiPSC-derived neurons infected with WNV NY99. Points show the mean of three independent replicates per time point; error bars indicate SD.

**Supplementary Fig. 3.**
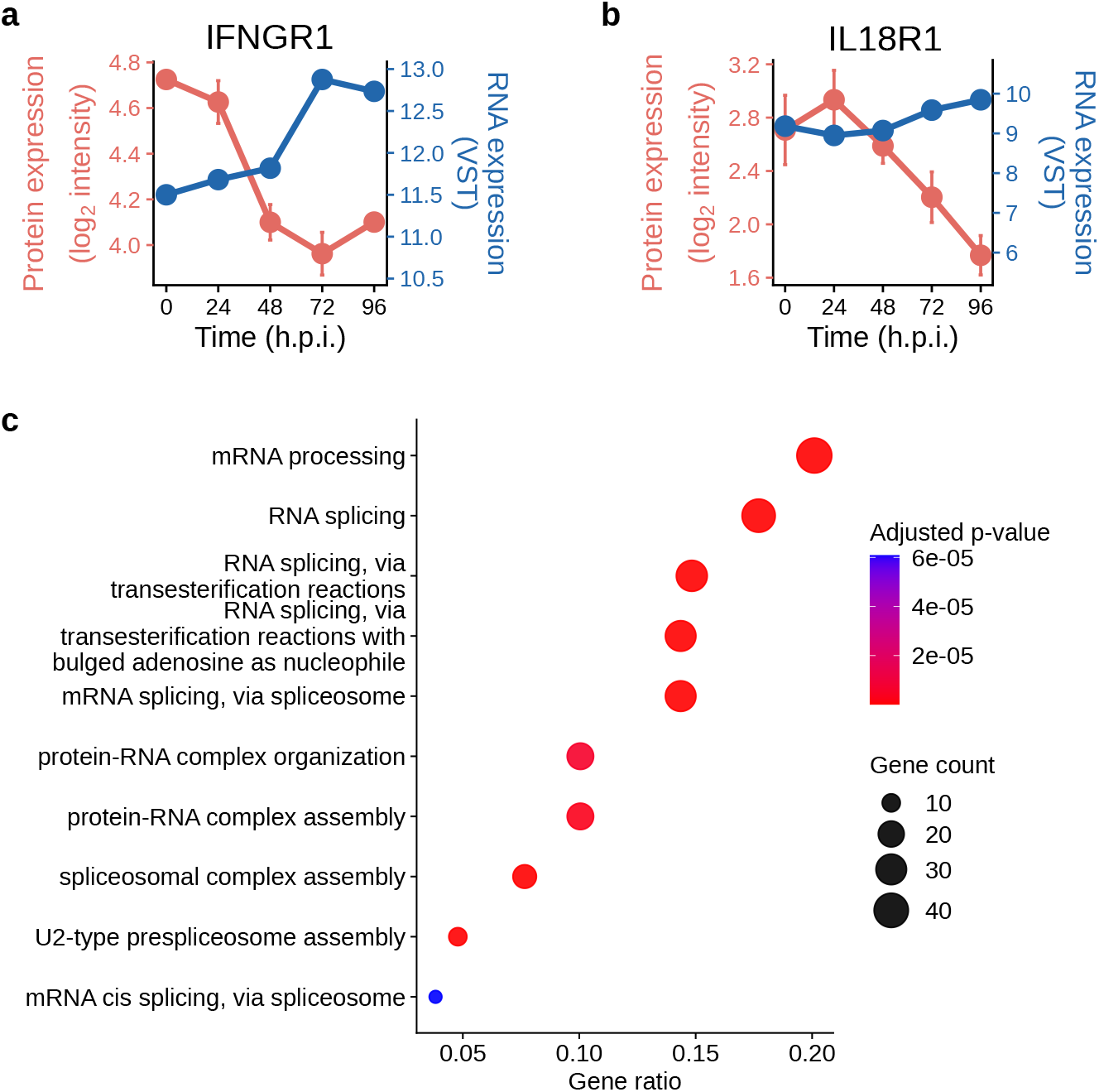
Protein and RNA levels are differentially regulated during infection. (a,b) Temporal changes in IFNGR1 (a) or IL18R1 (b) protein and RNA expression during WNV infection. (c) GO of biological processes enriched in the discordant set of proteins vs RNA abundance.

**Supplementary Fig. 4.**
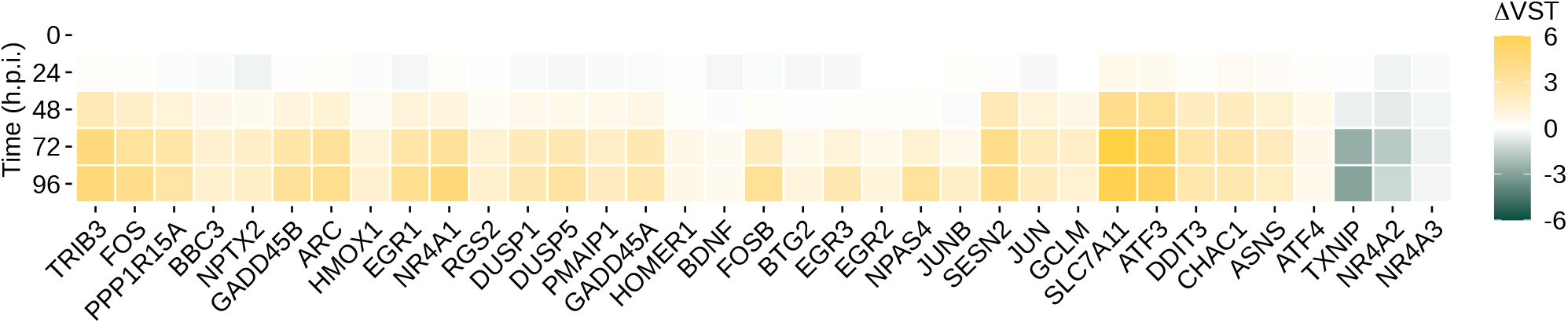
Temporal RNA expression profiles of selected neuronal activity-, calcium-responsive-, and cellular stress-associated genes during WNV infection. Heatmap showing changes in variance-stabilized transformed RNA expression (ΔVST) from 0 to 96 h.p.i., relative to the 0 h.p.i. baseline. Yellow indicates increased expression, green indicates decreased expression, and white indicates no change relative to baseline.

**Supplementary Fig. 5.**
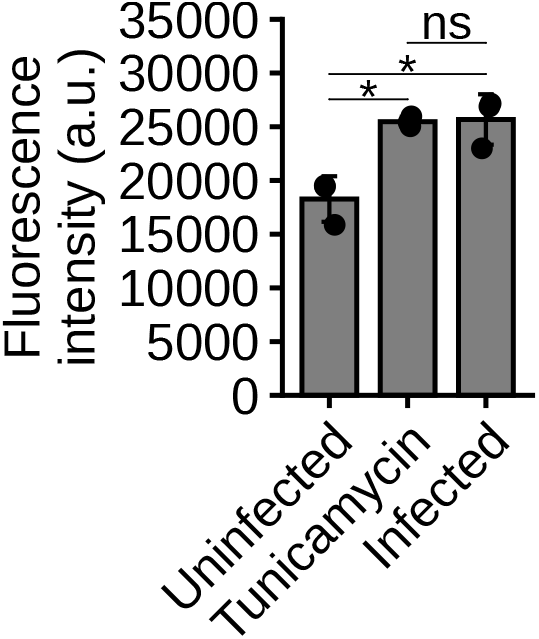
Calcium-dependent fluorescence intensity in uninfected, tunicamycin-treated, and WNV-infected neurons. Individual data points represent three independent differentiation replicates per condition. ns, not significant; *P < 0.05 (two-tailed unpaired Welch’s t-tests).

